# Disentangling the influences of pre- and postnatal periods on human cortical microstructure

**DOI:** 10.1101/2025.08.12.669812

**Authors:** Thanos Tsigaras, Juergen Dukart, Stuart Oldham, Simon B Eickhoff, Casey Paquola

## Abstract

During late gestation and early postnatal development a combination of intrinsic and extrinsic factors drive the maturation of the human cortex. This process is regionally heterogeneous, with cortical areas developing at different paces and trajectories. Leveraging submillimetre T1-weighted/T2w-weighted (T1w/T2w) magnetic resonance imaging (MRI) from pre- and full-term neonates (n = 599, 0-7 weeks), we sampled intracortical microstructure profiles across the cortex and characterised the profiles’ shapes according to their central moments. We found that gestational age at birth dominated the effects on early cortical development, with significant, global increases in microstructural density, increasing intracortical homogeneity and a bimodal change of the microstructural balance between superficial and deeper cortical layers. On the other hand, weeks since birth (i.e. postnatal age) exhibited different effects on microstructure, with density increasing at a slower pace, increasing intracortical heterogeneity, and intracortical balance only shifting towards deeper layers in posterior temporal, occipital, medial parietal areas and some prefrontal areas. These effects align with low spatial-frequency geometric eigenmodes of the human cortex, specifically the anterior-posterior, superior-inferior and central-polar axes. Our findings demonstrate that separating prenatal from postnatal influences, and analysing intracortical profiles rather than macroscale features, provides finer-grained insights into how human cortical microstructure changes during perinatal development and lays the groundwork for investigating the biological underpinnings that govern normative cortical maturation.

## Introduction

Perinatal neurodevelopment is a multifaceted procedure, comprising the early stages of the molecular, structural and functional maturation of the infantile human cortex. Birth marks the transition from the intrauterine, more genetically influenced development, to the extrauterine, increasingly sensory-driven development of the brain. *In utero* the foetus is relatively isolated from sensory input and therefore genetic and epigenetic programming primarily steer neurobiological processes, such as neuronal proliferation and migration (Bystron et al., 2008), axonal growth and synaptogenesis (Webb et al., 2001), in order to establish the cytoarchitectonic scaffold of the human cortex. After birth, aside from the intrinsic refinement of the cortical structure, sensory input drives neural patterning, with processes such as synaptic and axonal pruning, accompanied by myelination, refining and reorganising cortical connectivity (Huttenlocher & Dabholkar, 1997; Stiles & Jernigan, 2010). This intra-versus extrauterine dichotomy should be taken into consideration when interpreting changes in the cortical structure of the neonatal human brain, emphasising its lasting impact on the architecture of the developing brain.

Throughout gestation, multiple morphogenic gradients (e.g. FGF, Shh and WNT) define the general areal patterning of the human cortex (Borello & Pierani, 2010). These gradients result in spatially differential expression of transcription factors, which in turn regulate the regionalisation of cortical areas (Borello & Pierani, 2010). During the early gestational period, network activity is largely dominated by intrinsic subplate bursts (Mukherjee & Kanold, 2022), but at later stages, thalamocortical projections begin to relay weak sensory drive, slowly shifting toward more sensory-driven activity. After birth, several biological processes co-occur in the cortex, which result in the reorganisation and fine-tuning of the cortical microstructure (Brody et al., 1987; Huttenlocher & Dabholkar, 1997; Stiles & Jernigan, 2010). Key transitions that directly influence infantile neurodevelopment are the exchange of placental support for pulmonary respiration, the burst of visual sensory signals, changes in intracranial pressure and several metabolic switches. These events transfer developmental control from genetic preprogramming to increasing experience-dependent plasticity. Cortical connections that are used more frequently are reinforced via myelination and sparsely used axons or synapses are pruned. An interplay of thousands of molecules regulates these processes (Breen et al., 2018), rendering this period sensitive to both genetic and environmental factors that can underpin the onset of various neurodevelopmental disorders (Roth & Sweatt, 2011). Premature birth can also disrupt the intrinsic, primarily genetically guided neurodevelopmental processes, forcing them to proceed earlier in the extrauterine environment, outside of the protected intrauterine milieu. This directly affects subsequent development and is linked with long-term cognitive vulnerability (Gozdas et al., 2018; Smyser et al., 2016).

The process of early cortical maturation is characterised by heterochronicity, in which cortical regions develop at different paces. For example, synaptic density in the primary visual cortex peaks during the 5^th^ month of development, whereas in the prefrontal cortex, the peak occurs during the 15^th^ postnatal month (Huttenlocher & Dabholkar, 1997). Magnetic resonance imaging (MRI) measurements have also shown regionally variable patterns of cortical maturation (Ball et al., 2020; Li et al., 2013; Wang et al., 2019). Across multiple modalities, differences in cortical development are most prominent between sensory and association cortices, suggesting that unimodal areas develop earlier than heteromodal areas (Gao et al., 2015; Gilmore et al., 2012; Lyall et al., 2015). Complementing tissue-level findings, connectomic analyses suggest that by term age, the topology of structural hubs is distributed in an adult-like configuration across sensorimotor and association cortices, whereas functional hubs are initially concentrated in primary regions and migrate toward heteromodal areas during childhood (Oldham & Fornito, 2019).

Studies investigating the developing structure of the human cortex *in vivo*, have typically been constrained to macroscale features, such as thickness and surface area. Such features do not relate to cytoarchitectural changes (la Fougere et al., 2011; Ribeiro et al., 2013), that are fundamental to the formation of functional hierarchies in the cortex (Garcia-Cabezas et al., 2019). Histological studies, that do directly assess cytoarchitectural features, are naturally cross-sectional and often very limited in sample size. Hence, to better understand the early structural changes of the cortex, there is a need for non-invasive methods which can characterise cortical microstructure. T1-weighted (T1w) and T2-weighted (T2w) MR images can help to address this need as their ratio minimises the impact of magnetic inhomogeneities and enhances the microstructural signal. While initially considered a “myelin-marker”, (Glasser et al., 2014; Glasser & Van Essen, 2011; Nakamura et al., 2017), additional studies have emphasised the contribution of other microstructural features (e.g. iron concentration and cellular density) to the T1w/T2w signal (Fukunaga et al., 2010; Preziosa et al., 2021).

Leveraging sub-millimetre resolution MRI, T1w/T2w imaging can be used to investigate microstructural development across cortical depths (Paquola et al., 2019b; Sui et al., 2022). The laminar scaffold of the cortex varies across regions, with each area containing a characteristic stack of layers, where specific combinations of neurons receive or dispatch information streams (Harris & Shepherd, 2015). While the laminar and columnar organisation of the cortex is laid down during gestation, it continues to be intensively refined postnatally, a process that contributes to the emerging functional specialisation of each cortical region (Meyer et al., 2000; Molnar et al., 2019). MR-based intracortical profiling and histology-inspired central-moment metrics (Schleicher et al., 1999) have recently been used to reproducibly discriminate cytoarchitectonic areas (Amunts et al., 2020), track adolescent re-organisation of the sensory-fugal gradient (Paquola et al., 2019a), and chart the convergence of cortical types and functional motifs along the human mesiotemporal axis (Paquola et al., 2020). These applications enable the mapping of both region- and depth-specific maturational processes in an observer-independent manner.

In this study, we leveraged high-resolution MRI from 599 infants in their first postnatal weeks, spanning a wide spectrum of gestational ages (23 – 42 weeks), to characterise the maturational dynamics of cortical microstructure. We evaluate how intra- and extrauterine durations impact cortical development and its progression, as well as how modelling gestational and postnatal ages separately can help explain the regional differences in the influence of dominantly intrinsic, prenatal programming from increasingly experience-dependent postnatal changes on cortical microstructure. We finally assess to what extent these dynamics are constrained by low-dimensional cortical geometry.

## Methods

### Participants and demographics

The present study leveraged the Developing Human Connectome Project (dHCP), an open-access initiative designed to map neonatal brain development using multi-modal MRI [https://www.developingconnectome.org/, (Hughes et al., 2017)]. In the 4^th^ release of the dataset (https://biomedia.github.io/dHCP-release-notes/), 915 neonate participants (426 females, 26 – 45 weeks postmenstrual age) were provided, including term-born (n = 606) and prematurely born (n = 309) infants. Infants born at 37 weeks of gestational age or earlier are classified as preterm (Quinn et al., 2016). Exclusion criteria were chromosomal abnormalities, brain injuries, contraindications for MRI scanning and acute birth complications treated with prolonged resuscitation. Among the 915 participants, 100 originated from non-singleton pregnancies. Additionally, a subsample of neonates were scanned more than once, with the 4^th^ release including two timepoints for 106 participants and three timepoints for two participants.

Here, we examined the influence of three different developmental measures: gestational age, postnatal age, and postmenstrual age. Gestational age captures the intrauterine development of the foetus and is calculated as the duration from the first day of the mother’s last menstrual period to the day of birth. Postnatal age refers to the period from birth to the day of the scan, reflecting the extrauterine development of the infant. Finally, postmenstrual age is the sum of the former two metrics, representing the cumulative maturation of the brain during both the pre- and postnatal stages of development.

For our study, additional inclusion criteria were the availability of T1w and T2w images, as well as a complete FreeSurfer output. Furthermore, for participants with longitudinal scanning, we only used scans from the earliest timepoint to maintain a balanced cross-sectional dataset and avoid biases introduced by repeated measures. Finally, since the oldest term-born infant had a postnatal age of 6^+6^ weeks, scans of preterm neonates with a postnatal age exceeding 7 weeks were also excluded. The final dataset consisted of 599 participants (279 females, 162 preterm, 79 from non-singleton pregnancies).

Results were replicated using the same sample of participants but excluding the participants of non-singleton pregnancies (**Supp. Fig. 1-3**). Full demographics of the subsample used in this study are presented in **Table 1** and **Figure 1**.

**Table 1:** Dataset demographics and developmental metrics

| | Median ± SD<br>(range) | Sex<br>differences | Bivariate correlation<br>with PMA (r) | Bivariate correlation<br>with GA (r) | Bivariate correlation<br>with PNA (r) |
| --- | --- | --- | --- | --- | --- |
| Postmenstrual age<br>(PMA, weeks) | 40<sup>+3</sup><br>(26<sup>+5</sup> – 44<sup>+5</sup>) | t(597) = 0.29<br>p = 0.77 | - | 0.92 | -0.04 |
| Gestational age (GA,<br>weeks) | 39<sup>+2</sup><br>(23<sup>+5</sup> – 42<sup>+2</sup>) | t(597) = 0.10<br>p = 0.92 | 0.92 | - | -0.43 |
| Postnatal age (PNA,<br>weeks) | 0<sup>+6</sup><br>(0.00 – 7.00) | t(597) = 0.42<br>p = 0.68 | -0.04 | -0.43 | - |
| Weight at scan (kg) | 3.06 ± 0.92<br>(0.74 – 5.36) | t(559) = -2.25<br>p = 0.03 | 0.88 | 0.83 | -0.10 |
| Head circumference<br>at scan (cm) | 33.67 ± 3.14<br>(21.00 – 39.10) | t(571) = -2.44<br>p = 0.02 | 0.88 | 0.83 | -0.08 |

**Figure 1:**
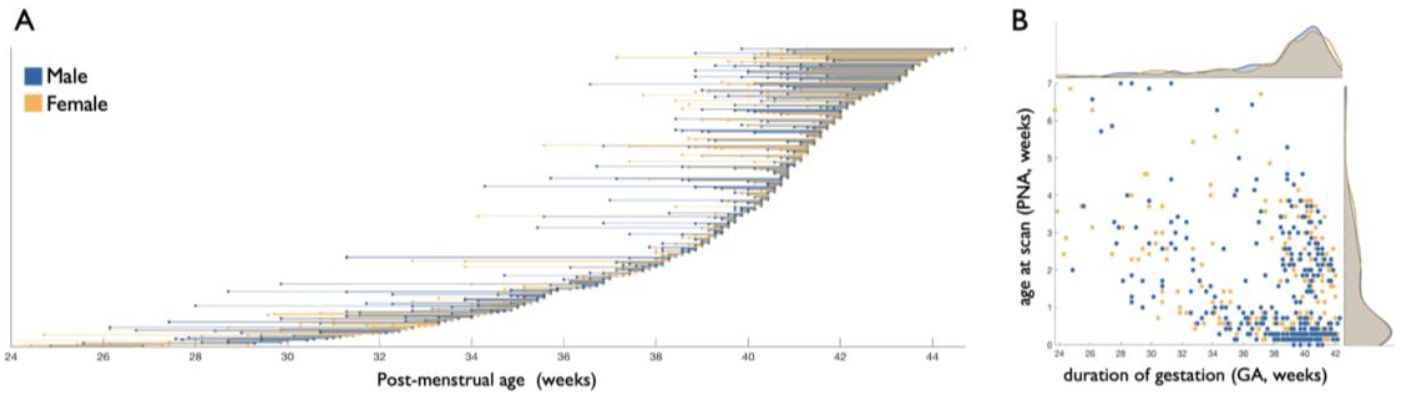
**A)** Age distributions of the 599 participants (y-axis). The plot represents duration of gestation at time of birth (left dot) and age at time of scan (right dot). **B)** The scatter plot illustrates the relationship between gestational age (GA) and postnatal age (PNA), with data points colour-coded by sex (blue: male, yellow: female). Histograms on the edges of the scatter plot show the distributions of gestational (top) and postnatal (right) age for males and females.

### MRI acquisition and preprocessing

The MRI acquisition was performed at St. Thomas Hospital, London using a Philips 3T scanner. The acquisition protocol was optimised for neonatal imaging, incorporating custom-built head coils as well as specialised positioning and immobilisation techniques to minimise motion artifacts, while ensuring the safety of the participants (Hughes et al., 2017). Participants were naturally asleep during the image acquisition with the exception of six out of the 805 subjects, for whom sedation was used (two of which included in the present study). T1 images were acquired using an IR (Inversion Recovery) TSE sequence at a resolution of 0.8 × 0.8 × 1.6 mm with a repetition time (TR) of 4.8s, an echo time (TE) of 8.7ms and sensitivity encoding (SENSE) factors of 2.26 (axial) and 2.66 (sagittal). T2 images were acquired at the same resolution using a Turbo Spin Echo (TSE) sequence, with a TR of 12s, TE of 156ms and SENSE factors of 2.11 (axial) and 2.58 (sagittal).

Preprocessed data were downloaded from the 4^th^ release of the dHCP. For comprehensive details on image preprocessing, refer to Makropoulos et al. (2018). In brief, motion correction and super-resolution reconstruction were employed on T1 and T2 images to achieve isotropic volumes with a resolution of 0.5mm^3^ (Cordero-Grande et al., 2018; Kuklisova-Murgasova et al., 2012). Then, brain tissue was extracted from the T2 images [BET, (Smith, 2002)], tissue-type segmentation was performed using Draw-EM (Makropoulos et al., 2014) and cortical surfaces were reconstructed from T1 images.

Following rigid co-registration of the motion-corrected T1w to the T2w image, T1w/T2w ratio images were computed. This method, introduced by Glasser and Van Essen (2011), minimises the impact of the inhomogeneous receive bias field and enhances the contrast between the grey matter (GM) and white matter (WM). The T1w/T2w ratio was initially interpreted as a proxy for myelin, as ratio maps aligned with known myeloarchitectonic borders and generally exhibited higher signal in areas with high myelin (e.g. primary sensorimotor cortex) (Glasser et al., 2016; Glasser & Van Essen, 2011). Additionally, significant differences in T1w/T2w values have been found between demyelinated and myelinated cortices of multiple-sclerosis patients (Nakamura et al., 2017) and on a transcriptomic level, higher levels of myelin-related transcripts are expressed in cortical areas with higher T1w/T2w values (Ritchie et al., 2018). Nevertheless, recent studies have questioned the specificity of the T1w/T2w ratio to myelin and have suggested that neurite density, iron concentration and other microstructural factors also have significant influence on intracortical variation in the T1w/T2w ratio (Righart et al., 2017; Sandrone et al., 2023). Taking all these findings into consideration, here we use intracortical T1w/T2w signals as a composite marker of cortical microstructure.

To minimise potential inaccuracies of vertex-to-vertex correspondence, we parcellated cortical measurements into broader regions using the von Economo atlas, which captures regional differences in laminar organisation (Scholtens et al., 2018; von Economo & Koskinas, 1925). We aligned the parcellation scheme to native cortical surfaces via the dhcpSym40 40-week (postmenstrual age) infant template surface (Williams et al., 2023). For this purpose, we used template-to-native registration spheres that were released with dHCP and multimodal surface matching (Robinson et al., 2018), optimised for alignment of sulcal depth. Regions within the limbic lobe (von Economo areas L_A1_, L_A2_, L_C1_, L_C2_, L_C3_, L_D_ and L_E_) were excluded from the analysis due to limited cortical thickness in these regions, which in turn reduced the capacity of intracortical intensity sampling. This omission resulted to 66 parcels (33 in each hemisphere). For validation of the results, all analyses were replicated using the Schaefer-200 atlas (Schaefer et al., 2018) (**Supp. Fig. 4-5**).

### Microstructure profiles and their central moments

To analyse the cortical microstructure of the participants, we constructed 12 equivolumetric surfaces between native pial and white matter surfaces. The equivolumetric model compensates for variations in laminar thickness between sulci and gyri (Bok, 1929; Consolini et al., 2022) by varying the Euclidean distance between pairs of intracortical surfaces based on curvature to preserve the fractional volume between surfaces (Waehnert et al., 2014). Then, we sampled image intensities at matched vertices across the 12 intracortical depths. (**Figure 2A**).

**Figure 2:**
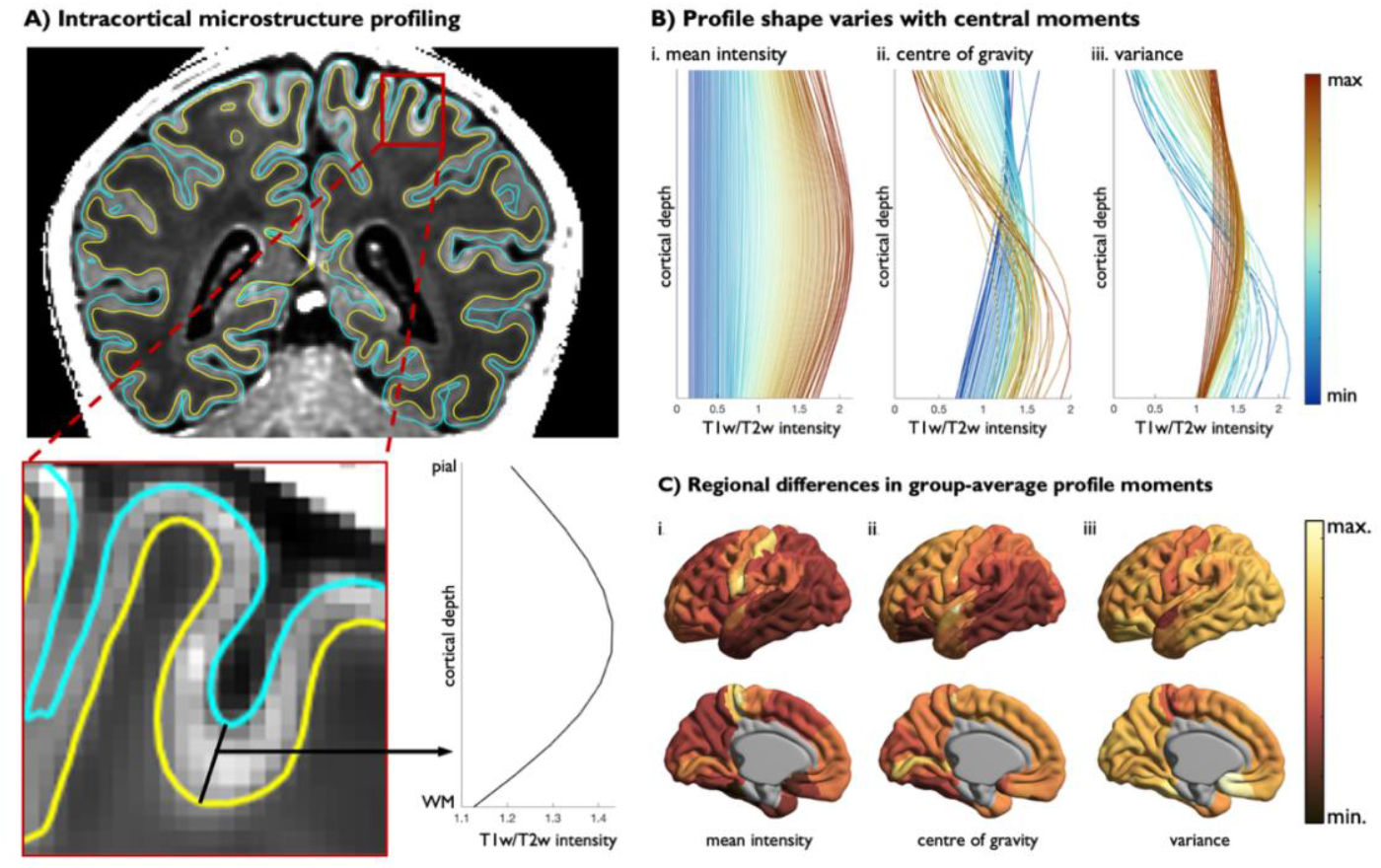
**A)** Intracortical intensity profiles were extracted at each vertex of infant structural MRI scans. Intensities were sampled at twelve equivolumetric intracortical surfaces, spanning from the pial boundary (blue) to the white matter boundary (yellow), capturing signal variations across cortical depths, defined as microstructure profiles. **B)** The shape of the average microstructure profile within each von Economo parcel was summarised using three central moments: mean intensity, centre of gravity, and variance. The mean intensity describes the average amplitude of the T1w/T2w signal across the profile, the centre of gravity indicates the relative depth at which the signal is concentrated, and the variance describes the dispersion of the intensity distribution across intracortical depths. **C)** Parcel-wise central moment distributions mapped on the dHCP 40-week surface template. Excluded regions (i.e. von Economo areas L_A1_, L_A2_, L_C1_, L_C2_, L_C3_, L_D_ and L_E_ and the cortical wall) are shown in grey.

To achieve a low-dimensional representation of the complex patterns of intracortical variation, we characterised the shape of each profile using three central moments. This method, inspired by histological studies (Schleicher et al., 1999), provides a compact, biological interpretable description of the signal distributions. The zeroth moment, *mean intensity (μ*_*0*_), represents the average signal amplitude across intracortical depths (**Figure 2Bi**). The first moment, *centre of gravity (μ*_*1*_), describes how the intensity distribution is balanced across cortical depths, whereby a high centre of gravity corresponds to higher intensities in the deeper cortex and a low centre of gravity describes higher intensities in the superficial cortex, hence providing an indicator of the balance of microstructure across supra- and infragranular layers (**Figure 2Bii**). Finally, the second moment, *variance (μ*_*2*_), characterises the spread of intensities across the profile. A higher variance relates to a flatter intensity profile (i.e. uniformly distributed intensities across intracortical depths) and a low variance describes a heterogeneous intensity distribution across intracortical depths (**Figure 2Biii**). Each of these moments, as well as their combinations, provide insights to the degree of laminar differentiation in the cortex and capture distinct patterns of microstructural differentiation across the cortical surface (**Figure 2C**).

### Statistical analysis

The statistical analysis focused on investigating the effects of postmenstrual age, gestational age and postnatal age on microstructure, defined by the central moments of average intracortical intensity profiles on a parcel level. These three developmental metrics were analysed both independently and collectively to account for different aspects of cortical development. First, to assess cumulative maturation of the brain, parcel-wise linear regression analysis was performed using postmenstrual age and sex as predictors and each central moment individually as response variables. False discovery rate (FDR) correction with an alpha of 0.025 was applied to account for multiple comparisons.

Next, to dissect the effect of postmenstrual age on cortical microstructure into its pre- and postnatal components, we first performed a linear model comparison, with each average central moment across all parcels as the response variable and combinations of gestational age, postnatal age and sex as predictors, validating the fit of these variables for modelling cortical microstructure. Due to the broad overlap of the distributions of gestational and postnatal age, we isolated the unique contributions of each developmental variable to cortical microstructure in multi-variate regression models, that included gestational age, postnatal age and sex. Results of univariate models of gestational and postnatal age are shown in **Supp. Fig. 6**. To validate the independence of the results from cortical thickness maturation, all results were replicated using models which additionally included cortical thickness as a predictor (**Supp. Fig. 7-9**).

To test whether regional variation in development concords with large-scale spatial gradients, we computed the product-moment correlations between the postmenstrual, gestational and postnatal age effects and the 2^nd^-4^th^ geometric eigenmodes (Pang et al., 2023). These modes are intrinsic spatial patterns derived from the shape of the cortex and were generated on the dhcpSym40 40-week surface template. A 1^st^ eigenmode is spatially uniform by construction and was therefore not included in the analysis. Product-moment correlations were performed on unthresholded vertex-wise maps and spin permutation testing (Alexander-Bloch et al., 2018) was performed to assess the significance of the correlation with the eigenmodes, while accounting for spatial autocorrelation in each map (10000 permutations, alpha level of 0.025). In addition, we assessed the additive effects of eigenmodes in explaining age effects through a series of uni-then multi-variate linear regressions.

## Results

### Changes in cortical microstructure with postmenstrual age

Globally, intracortical microstructure increases with postmenstrual age (r_mean_ = 0.68, p<0.001), in a manner that generally leads to more balanced distribution of the T1w/T2w signal across cortical depths (r_variance_ = 0.51, p<0.001). Increases in mean were most prominent in frontal and temporal regions, while increases in profile variance were stronger in occipital and parietal regions (**Figure 3A**). These changes resulted from microstructural increases in deeper layers for superior regions and increases in upper layers in inferior regions, as illustrated by the bimodal pattern of age-related changes in centre of gravity (**Figure 3A,B**). These analyses show how perinatal development of intracortical microstructure is region- and depth-specific, whereby unique developmental trajectories are expressed across the cortex.

**Figure 3:**
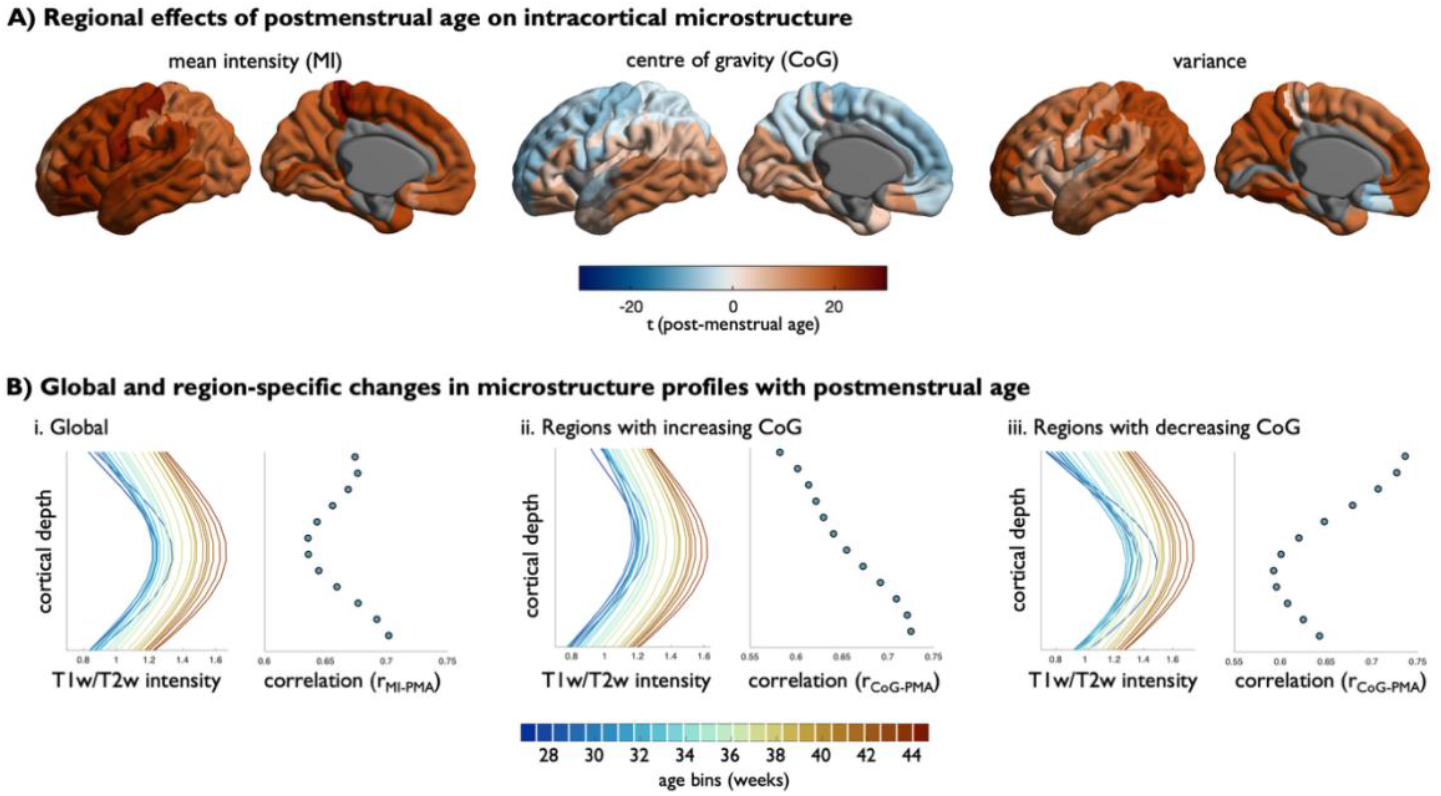
**A)** Linear regression models were used to assess the association between postmenstrual age (PMA) and intracortical profile moments across cortical regions, while controlling for sex. Surface maps display t-values for the PMA-estimate, projected onto the cortical surface for mean intensity (left), centre of gravity (middle) and variance (right). Statistically non-significant parcels (p > 0.05) and excluded parcels are displayed in grey. **B)** Sliding window visualisation of profile changes with PMA (age bins = 1.5-week windows with 50% overlap), with neighbouring scatter plots depicting the relationship between PMA with depth-specific T1w/T2w-intensities based on product-moment correlations. Abbreviations: MI: mean intensity; CoG: centre of gravity; PMA: postmenstrual age.

### Effects of gestational and postnatal age on cortical development

Next, we divided the developmental period into gestational age and postnatal age. To examine how the timing of birth and the developmental period after birth influence early cortical development, we linearly modelled the relationship of each central moment with gestational age and postnatal age, while controlling for the effect of the other (**Figure 4**). Gestational age effects mirrored the postmenstrual age effects reported above, emphasising the importance of the *in utero* period in shaping perinatal microstructure. During the postnatal period, we observed milder increases in mean microstructure, continuing in the same direction as during the *in utero* period. In contrast, we observed decreases in profile variance during the postnatal period, suggesting that cortical layers become more differentiable in the first postnatal weeks. Thus, while intracortical microstructure increases throughout the pre- and postnatal periods, its distribution across cortical layers shifts in distinct, region-specific ways in each period.

**Figure 4:**
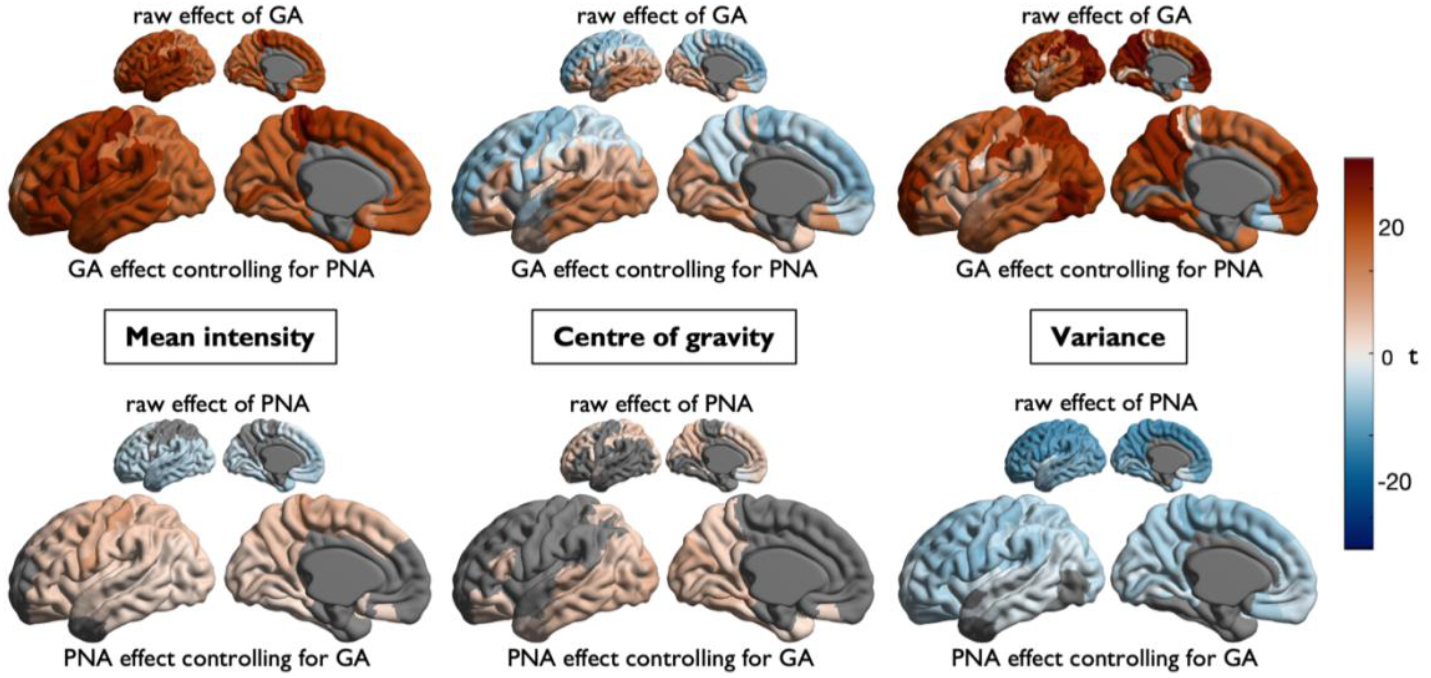
Large cortical surface maps illustrate gestational (GA, top) and postnatal (PNA, bottom) age effects on each central moment, after controlling for PNA and GA respectively. Smaller cortical surface maps illustrate univariate GA- and PNA-effects (i.e. without regressing out the other metric). Statistically non-significant parcels (p > 0.05) and excluded parcels are displayed in grey.

### Alignment of developmental effects on microstructure to geometric eigenmodes

Next, we assessed whether large-scale spatial gradients could account for regional differences in early microstructural development (**Figure 5**). Indeed, prenatally, changes in mean intensity were correlated with the 4^th^ eigenmode, capturing a central-polar axis (r = -0.48, p_spin_ = 0.02) and changes in centre of gravity were correlated with the 2^nd^ eigenmode, reflecting an anterior-posterior gradient (r = -0.62, p_spin_ = 0.03). Regional differences in postnatal age effects were also correlated with spatial axes, though with different axes to the prenatal effects. In particular, postnatal changes in mean intensity and variance aligned with the 3^rd^ eigenmode, capturing a superior-inferior axis (r_MI_ = -0.65, p_spin_ = 0.002; r_var_ = 0.57, p_spin_ = 0.04), while changes in the centre of gravity were aligned with the anterior-posterior axis (r = - 0.68, p_spin_ = 0.006). Combining eigenmodes in multi-variate models explained even more variance in the spatial patterns (**Figure 5B**). Age-related changes in centre of gravity were particularly strongly associated with eigenmodes (R^2^ > 0.6), suggesting that the preference of microstructural increases to occur in upper vs. lower layers is especially tightly linked to cortical geometry.

**Figure 5:**
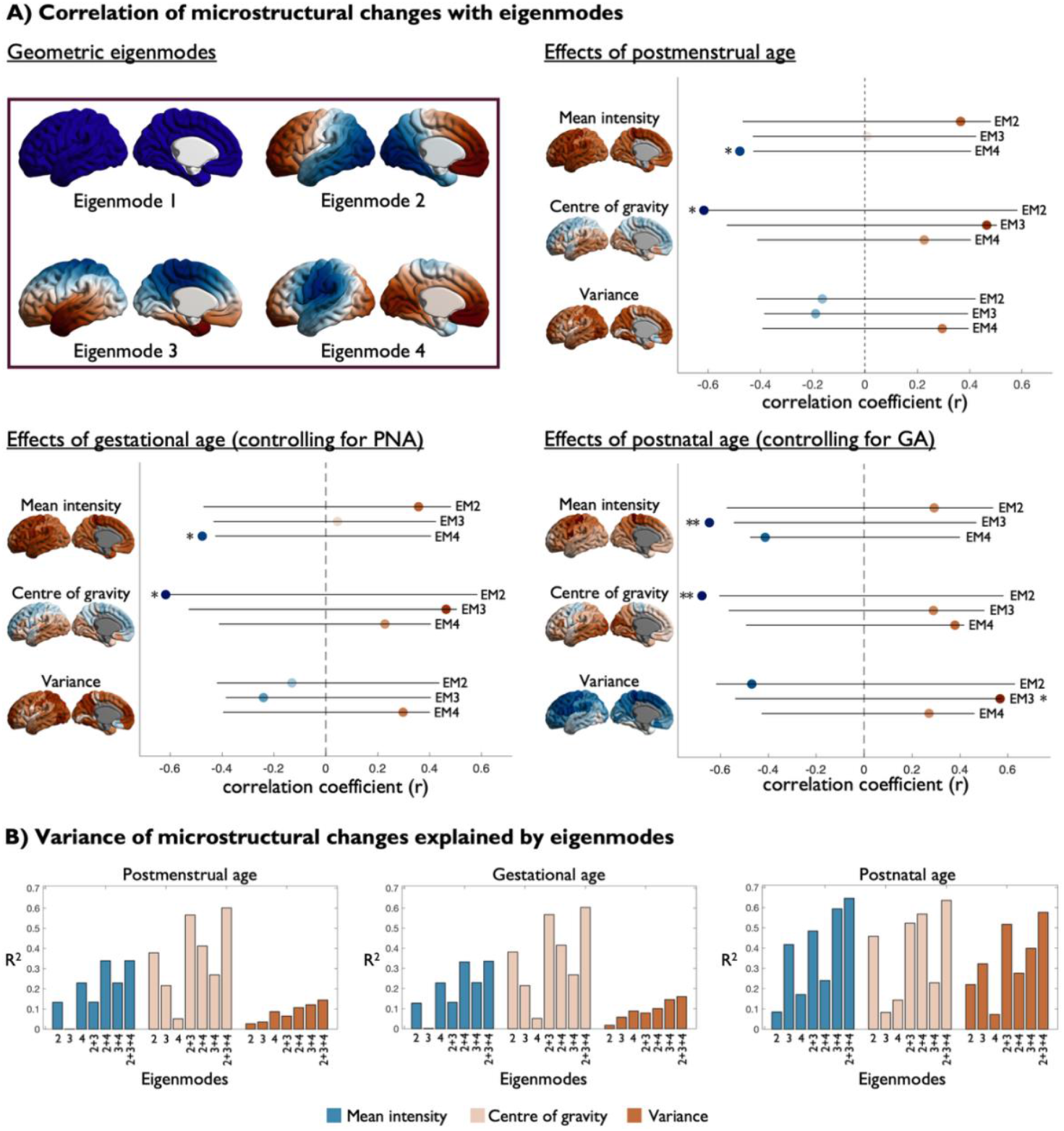
**A)** Product-moment correlation coefficients between the effects of postmenstrual (top right), gestational (bottom left) and postnatal age (bottom right) on each central moment and the 2^nd^-4^th^ eigenmodes. The whiskers represent the range of correlation values obtained from spin-based permutation testing (n = 10000), between the 2.5th to 97.5th percentile of the permuted correlations. Significant correlations are highlight with one asterisk for p < 0.05 and two asterisks for p < 0.01. **B)** Linear regression models were used to assess the explained variance of the effects of postmenstrual (left), gestational (middle) and postnatal age (right) by the 2^nd^-4^th^ geometric eigenmodes or combinations thereof.

## Discussion

Late gestation and early postnatal life are pivotal windows for the development of the human cortex, during which changes in cortical cytoarchitecture set the foundations for the brain’s functional specialisation. In this study, we combined histology-inspired intracortical profiling (Schleicher et al., 1999) with T1w/T2w ratio (Glasser & Van Essen, 2011) to characterise the developing structure of the infant human cortex. Using these methods, we identified regional heterogeneity in the structural changes that occur during the perinatal maturation of the cortex. Breaking down this perinatal period into its intra- and extrauterine stages revealed separable contributions of gestational and postnatal age to cortical development, highlighting how the structure of the cortex is differentially influenced by the internal environment during gestation and the external environment after birth. We also identified significant correlations between the developmental dynamics of microstructure and geometric eigenmodes of the cortex (Pang et al., 2023), suggesting that early cortical maturation is also shaped by topographical constraints.

The influences on cortical development shift at the time of birth. During intrauterine development intrinsic genetic programming primarily dictates the organisation of cortical laminae, and subplate activity guides neuronal migration and axonal extension (Cadwell et al., 2019; Kostovic et al., 2015). While sensory experience also shapes the cortical structure during gestation (Kostovic & Judas, 2010), after birth, the variety of sensory input and the maturation of GABAergic inhibition (Begum & Sng, 2017; Sale et al., 2010) shift the modulation of structural dynamics toward experience-dependent synaptogenesis and intracortical myelination (Huttenlocher & Dabholkar, 1997; Kuhne et al., 2021). In essence, perinatal development is characterised by a transition from a more strictly-regulated, genetically-governed maturation to experience-dependent changes in connection strength. This modulatory transition highlights the importance of the timing of birth in brain development, with prematurely born infants being exposed to a sensory-rich environment before the scaffold of their circuitry has been completely established for postnatal plasticity.

Addressing the transitional period around birth, our study highlighted the importance of investigating and characterising early cortical development in a way that assesses both the cumulative period of development (intra- and extrauterine), as well as each of its components individually. In particular, we included pre-term infants in our analyses to gauge a broad range of gestational durations and statistically compensated for the generally higher post-natal ages in pre-term infants using multi-variate regression models. By modelling gestational and postnatal ages separately, we showed that the timing of birth dominated the effects on early microstructural development, highlighting the importance of the intrinsic programming. We also observed significant postnatal changes in several cortical regions, even after controlling for these prominent effects of gestational maturation and despite the short postnatal period investigated in this study. Microstructural density continues to increase during early extrauterine development, but seemingly at a slower rate than during gestation. In occipital, posterior temporal, and some prefrontal regions, microstructure increases are more prominent in the deeper cortical layers, following a similar trajectory to the gestational dynamics. In contrast, we did not observe a continuation of prenatal increases in upper cortical layers in frontal and central areas in the postnatal period, showing a potential discontinuity between developmental trajectories of the two periods. Finally, whereas intracortical profiles become increasingly balanced with later gestational age, this effect seemed to flip after birth with postnatal changes in laminar organisation. In general, the prominent effect of gestational maturation on cortical microstructure emphasises the importance of the timing of birth and indicates that premature transition to the *ex utero* environment could disturb the well-regulated developmental trajectories of cortical growth, an insight that aligns with literature across various modalities (Ball et al., 2013; Dimitrova et al., 2021; Fenn-Moltu et al., 2023; Franca et al., 2024). Longitudinal studies extending into later infancy would be especially helpful for elucidating how and when the modest postnatal age influences observed here evolve into larger, experience-driven changes. Furthermore, such work could help to compensate for the imbalance in post-natal ages between pre-term and full-term infants and provide insight into how pre-term infants catch up (or don’t) on neurodevelopmental milestones (Bucher et al., 2002).

Regional differences in cortical microstructure originate from morphogen gradients, which pattern the mammalian cortex with variable concentrations of growth factors during gestation (Sansom & Livesey, 2009). Supporting the importance of gradients in shaping cortical development, we also found that large-scale eigenmodes capture a majority of variance in regional heterogeneity, especially in terms of changes in the balance between upper and lower cortical layers. During postnatal development, thalamocortical connectivity is thought to refine cortical arealisation, especially around primary sensory areas (Cadwell et al., 2019; Molnar & Kwan, 2024). Relatedly, histological and imaging studies consistently show that primary sensory and motor regions mature earlier than association cortex (Li et al., 2013; Natu et al., 2021; Wang et al., 2019). While the short postnatal timeframe encompassed by this dataset precludes interpretation of maturity per se, we observed strong postnatal changes in upper layers of the occipital and inferior temporal cortex, highlighting a preferential development of upper layers of cortical regions involved in visual processing during this early period of immense sensory stimulation. Notably, we found an opposing trend in sensorimotor areas, with a preference towards microstructural development of deeper layers. This contrast may underpin the differentiation of sensorimotor and visual cortical areas; a pattern that aligns with their strong functional differentiation during infancy (Lariviere et al., 2020).

In conclusion, the present study demonstrates how translating intracortical microstructure profiling can reveal distinct spatial patterns of development, which reflect the regional differentiation across large-scale spatial gradients. These findings offer a framework for linking macroscale in-vivo MRI to microscale biology, showing how depth-specific changes underpin regional differentiation between sensory systems and levels of the cortical hierarchy. Modelling gestational and postnatal ages separately helped to disentangle intrinsic, prenatal influences from more experience-dependent postnatal changes of cortical microstructure, illustrating how intrauterine development dominates microstructural refinement in comparison to the very early stages of extrauterine development. Our approach lays the groundwork for future studies to elucidate the biological underpinnings that govern normative cortical maturation, as well as to pinpoint how prematurity increases the risk of neurodevelopmental disorders (Crump et al., 2021, 2023; Franz et al., 2018).

## Acknowledgements

This work was supported by the Deutsche Forschungsgemeinschaft (DFG) under the Emmy Noether Programme (Project number - 524408221) and the German Scholars Organisation through the Klaus Tschira Boost Fund (KT40).

