## Supplementary for "Disentangling the influences of pre- and postnatal periods on human cortical microstructure"

### Effects of postmenstrual age on cortical microstructure, excluding twin participants

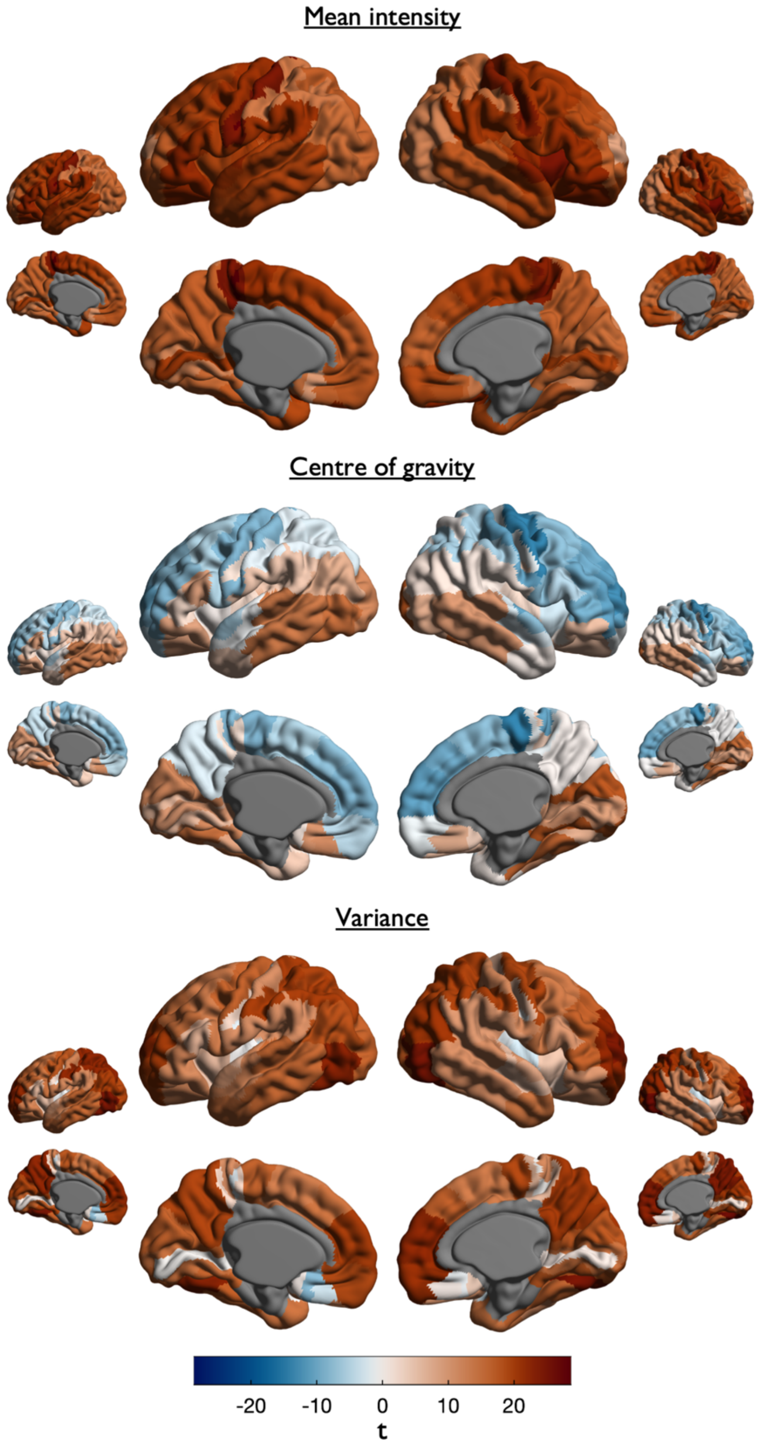


**Supplementary figure 1:** Linear regression models were used to assess the association between postmenstrual age (PMA) and intracortical profile moments across cortical regions, while controlling for sex. Bigger surface maps were derived from a subsample of participants, which excluded any participants of multiple pregnancies. Smaller surface maps were derived from the original sample of this study. Surface maps display t-values for the PMA-estimate, projected onto the cortical surface for mean intensity (top), centre of gravity (middle) and variance (bottom). Excluded parcels are displayed in grey.

### Effects of gestational age on cortical microstructure, excluding twin participants

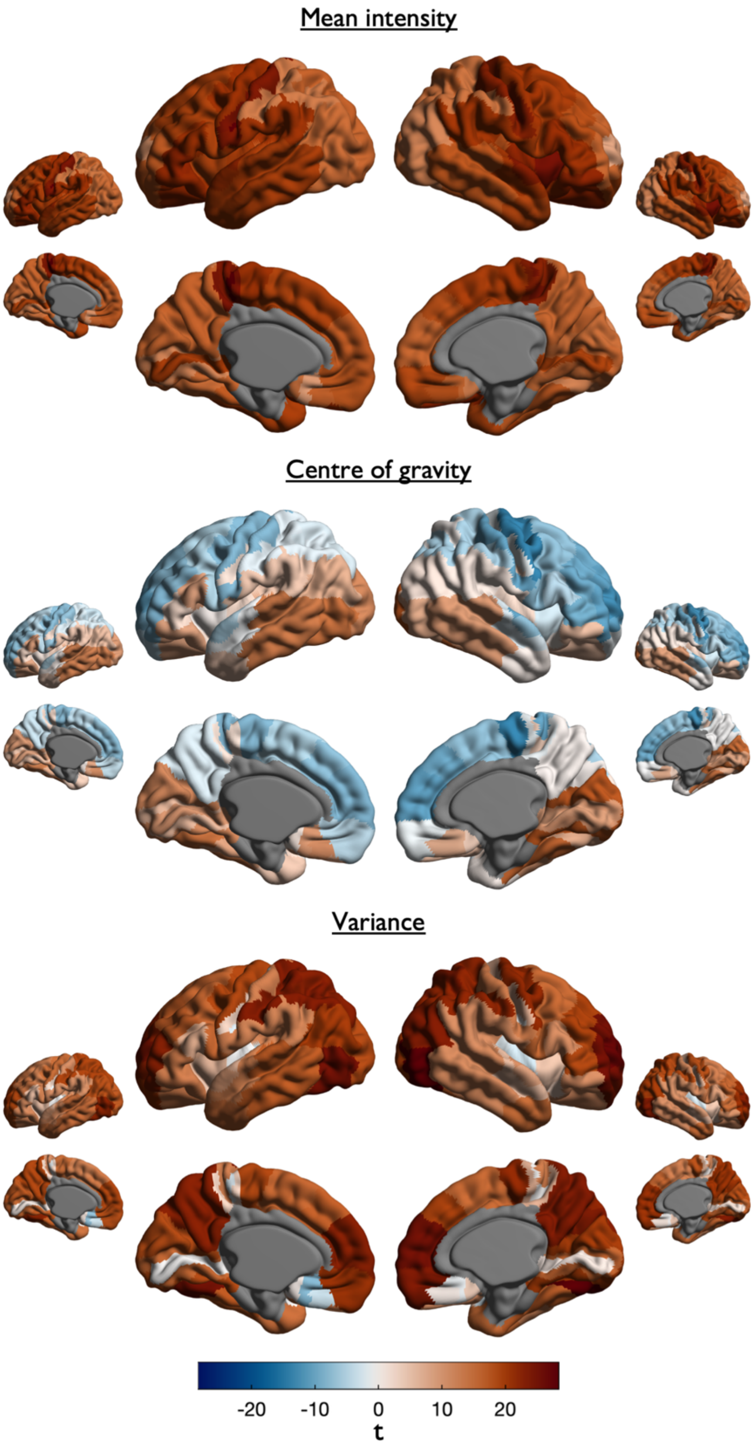


**Supplementary figure 2:** Linear regression models were used to assess the association between gestational age (GA) and intracortical profile moments across cortical regions, while controlling for sex. Bigger surface maps were derived from a subsample of participants, which excluded any participants of multiple pregnancies. Smaller surface maps were derived from the original sample of this study. Surface maps display t-values for the GA-estimate, projected onto the cortical surface for mean intensity (top), centre of gravity (middle) and variance (bottom). Excluded parcels are displayed in grey.

### Effects of postnatal age on cortical microstructure, excluding twin participants

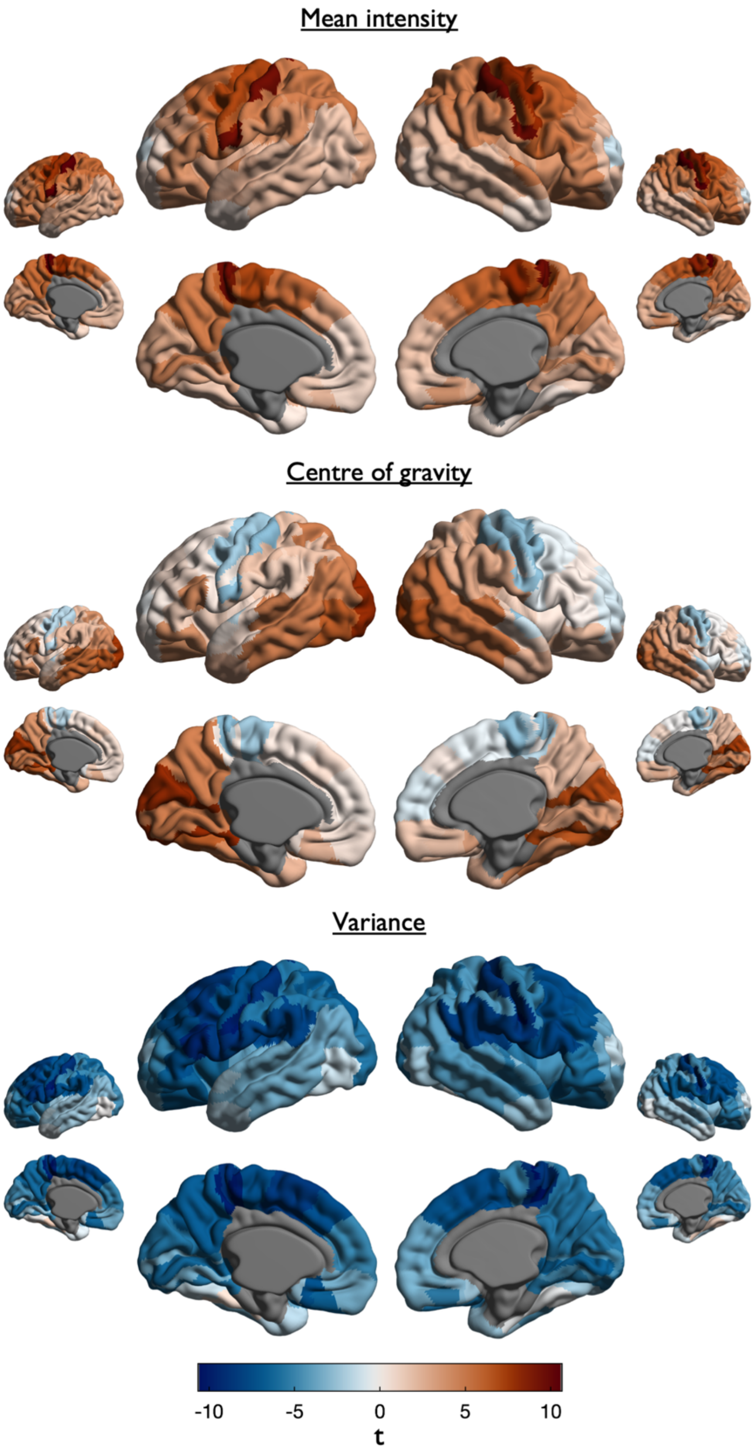


**Supplementary figure 3:** Linear regression models were used to assess the association between postnatal age (PNA) and intracortical profile moments across cortical regions, while controlling for sex. Bigger surface maps were derived from a subsample of participants, which excluded any participants of multiple pregnancies. Smaller surface maps were derived from the original sample of this study. Surface maps display t-values for the PNA-estimate, projected onto the cortical surface for mean intensity (top), centre of gravity (middle) and variance (bottom). Excluded parcels are displayed in grey.

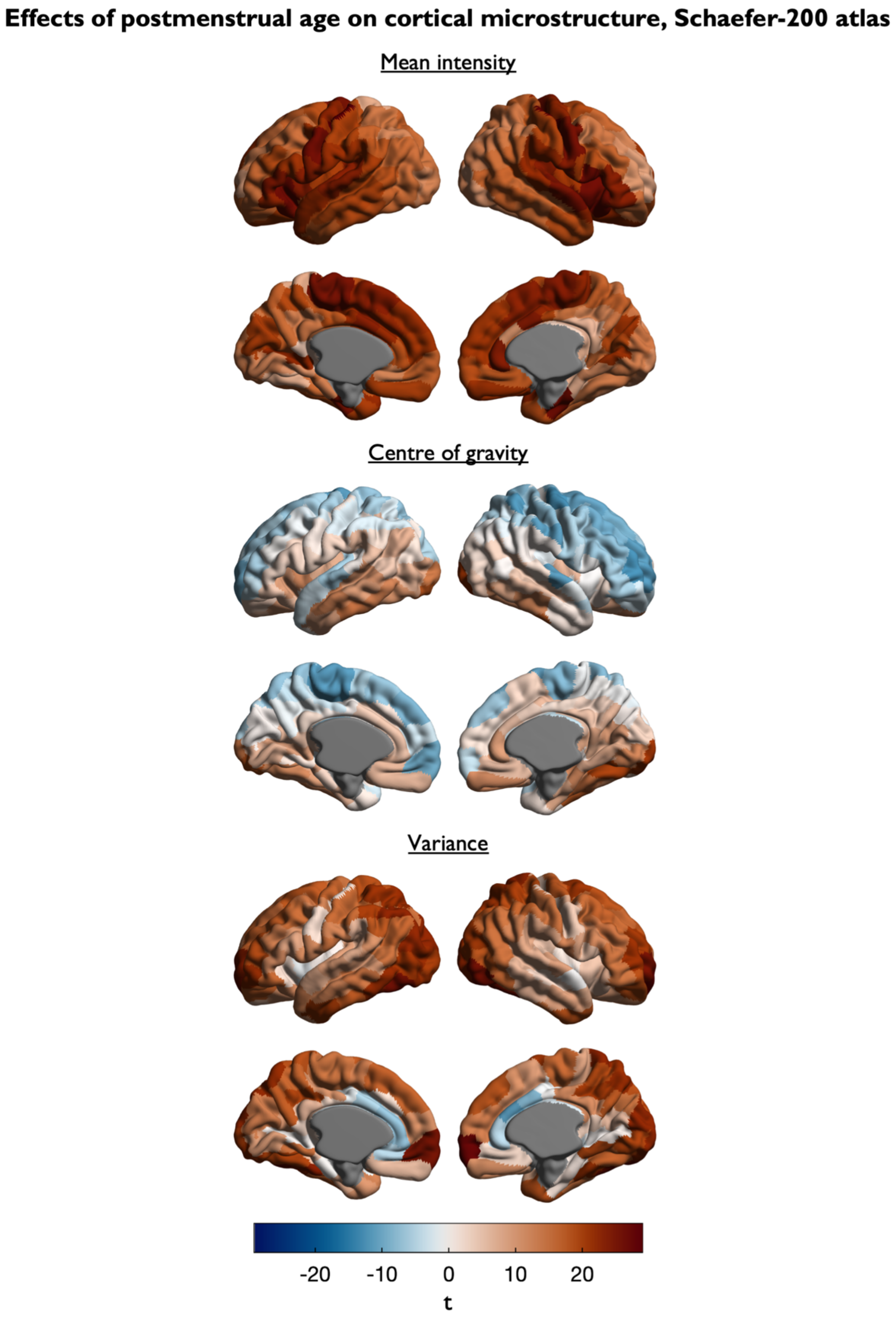


**Supplementary figure 4:** Linear regression models were used to assess the association between postmenstrual age (PMA) and intracortical profile moments across cortical regions of the Schaefer-200 atlas, while controlling for sex. Surface maps display t-values for the PMA-estimate, projected onto the cortical surface for mean intensity (top), centre of gravity (middle) and variance (bottom). Excluded parcels are displayed in grey.

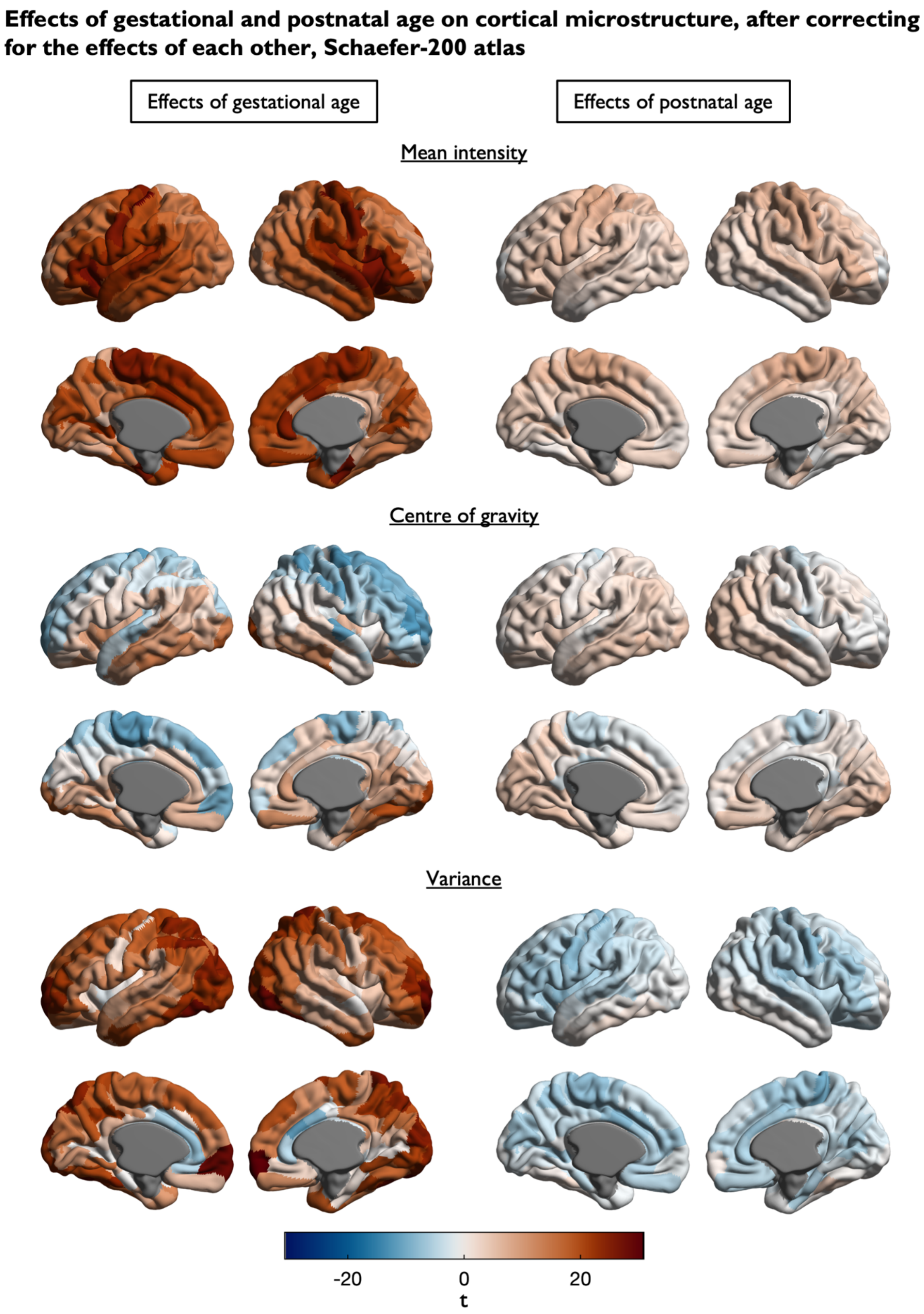


**Supplementary figure 5:** Linear regression models were used to assess the association between gestational age (GA, left) and postnatal age (PNA, right) with intracortical profile moments across cortical regions of the Schaefer-200 atlas, while controlling for sex. Surface maps display t-values for the GA- and PNA-estimates, projected onto the cortical surface for mean intensity (top), centre of gravity (middle) and variance (bottom). Excluded parcels are displayed in grey.

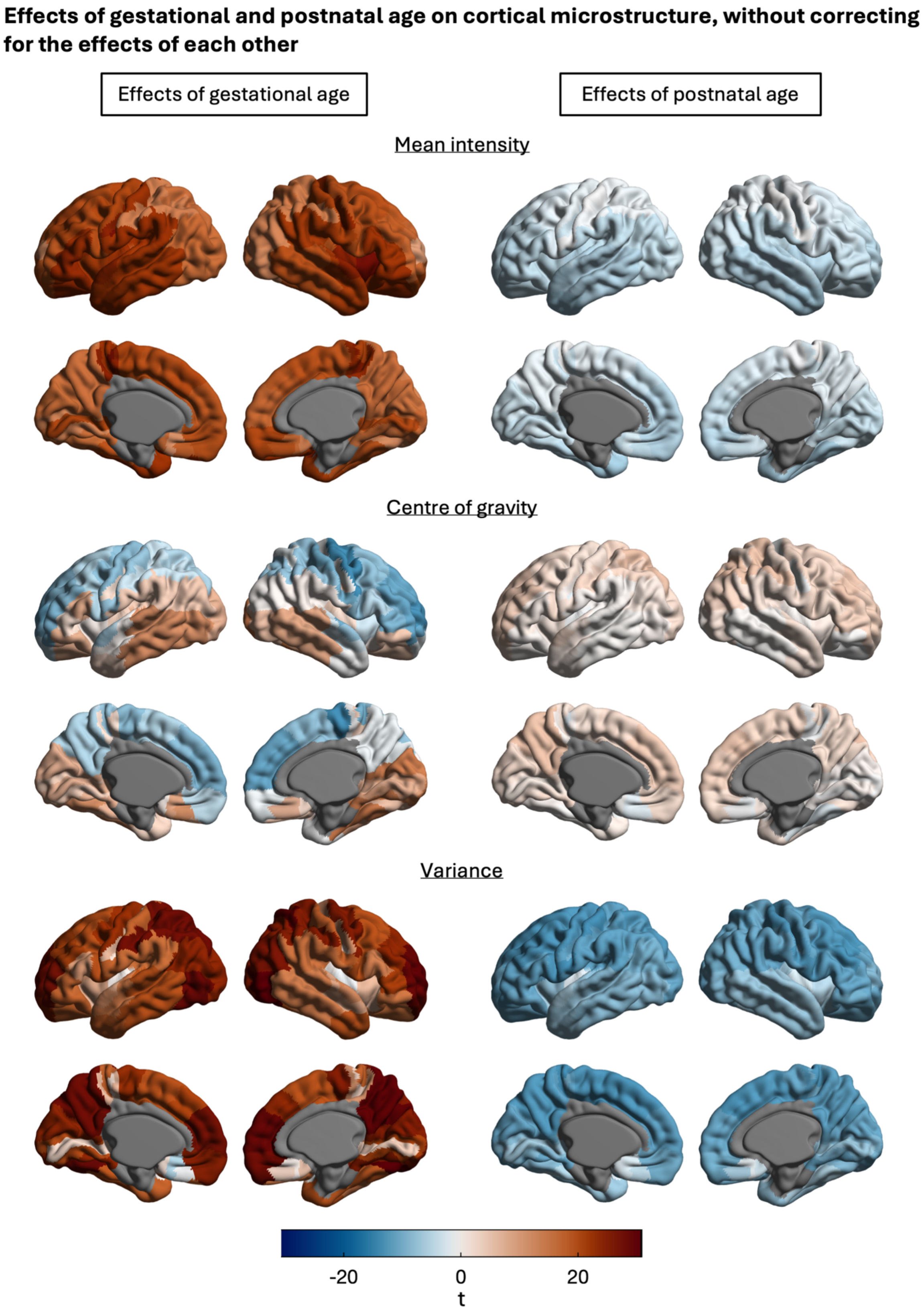


**Supplementary figure 6:** Cortical surface maps illustrate the spatial distribution of gestational (left) and postnatal (right) age effects on each central moment, without controlling for postnatal and gestational age respectively. The parcel-wise estimates were obtained from linear regression models with gestational age and sex or postnatal age and sex as predictors. Excluded parcels are displayed in grey.

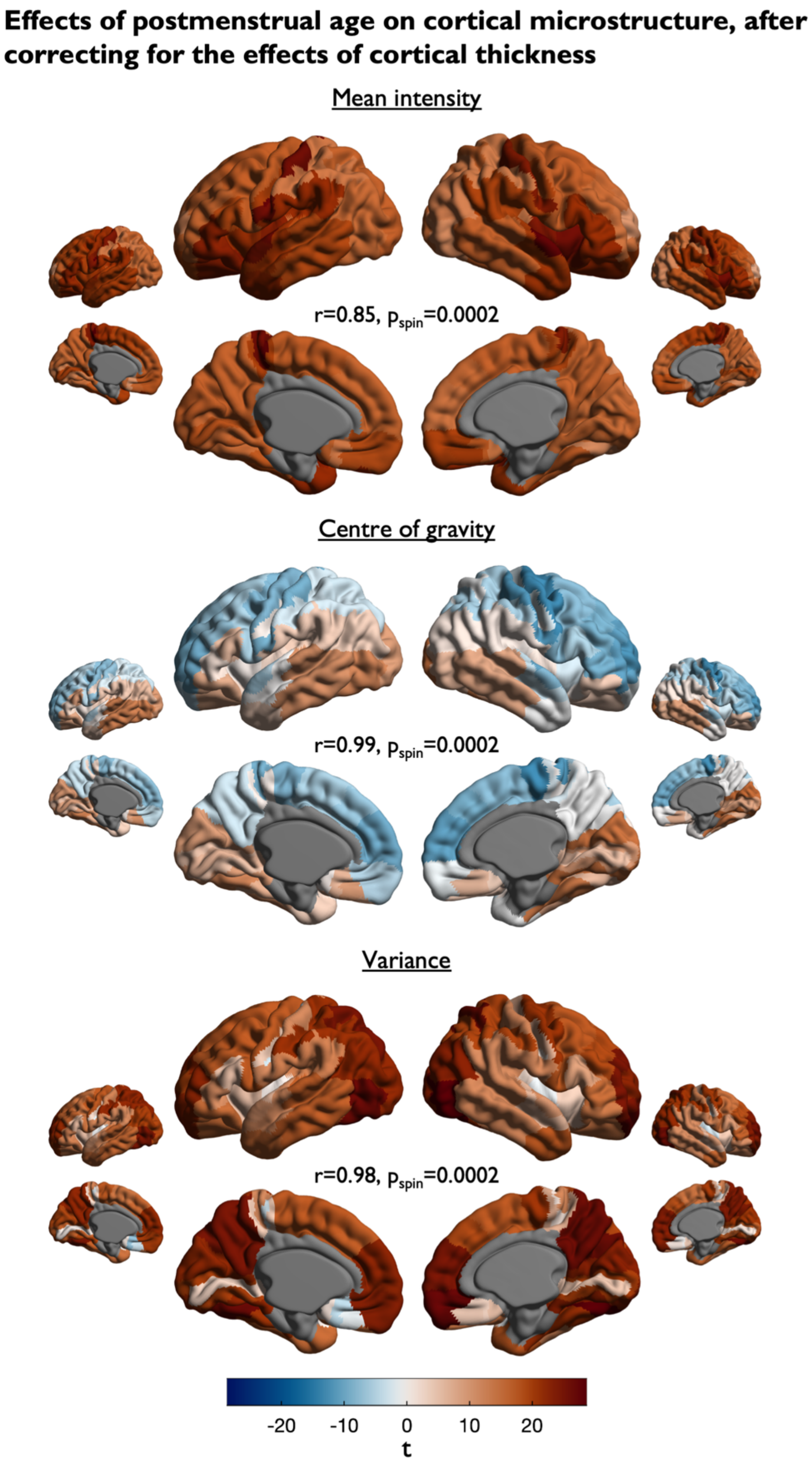


**Supplementary figure 7:** Linear regression models were used to assess the association between postmenstrual age (PMA) and intracortical profile moments across cortical regions, while controlling for cortical thickness and sex. The original results of this study, without controlling for cortical thickness are displayed as smaller surface maps. The spatial correlation between the effects with and without controlling for cortical thickness is displayed in the middle of each set, together with the p-value derived from spin-based permutation testing (n = 10000), between the 2.5th to 97.5th percentile of the permuted correlations. Surface maps display t-values for the PMA-estimate, projected onto the cortical surface for mean intensity (top), centre of gravity (middle) and variance (bottom). Excluded parcels are displayed in grey.

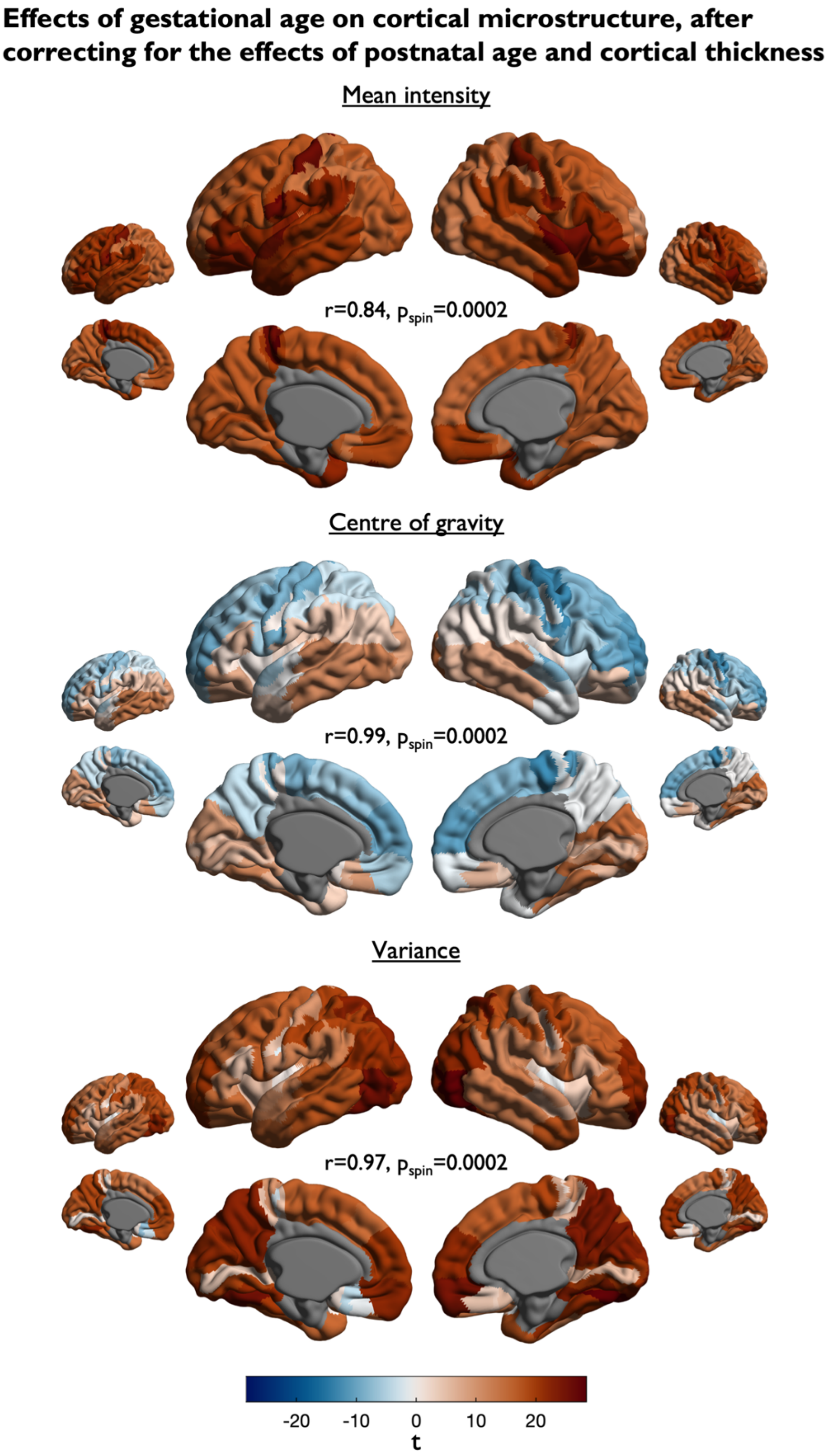


**Supplementary figure 8:** Linear regression models were used to assess the association between gestational age (GA) and intracortical profile moments across cortical regions, while controlling for cortical thickness and sex. The original results of this study, without controlling for cortical thickness are displayed as smaller surface maps. The spatial correlation between the effects with and without controlling for cortical thickness is displayed in the middle of each set, together with the p-value derived from spin-based permutation testing (n = 10000), between the 2.5th to 97.5th percentile of the permuted correlations. Surface maps display t-values for the GA-estimate, projected onto the cortical surface for mean intensity (top), centre of gravity (middle) and variance (bottom). Excluded parcels are displayed in grey.

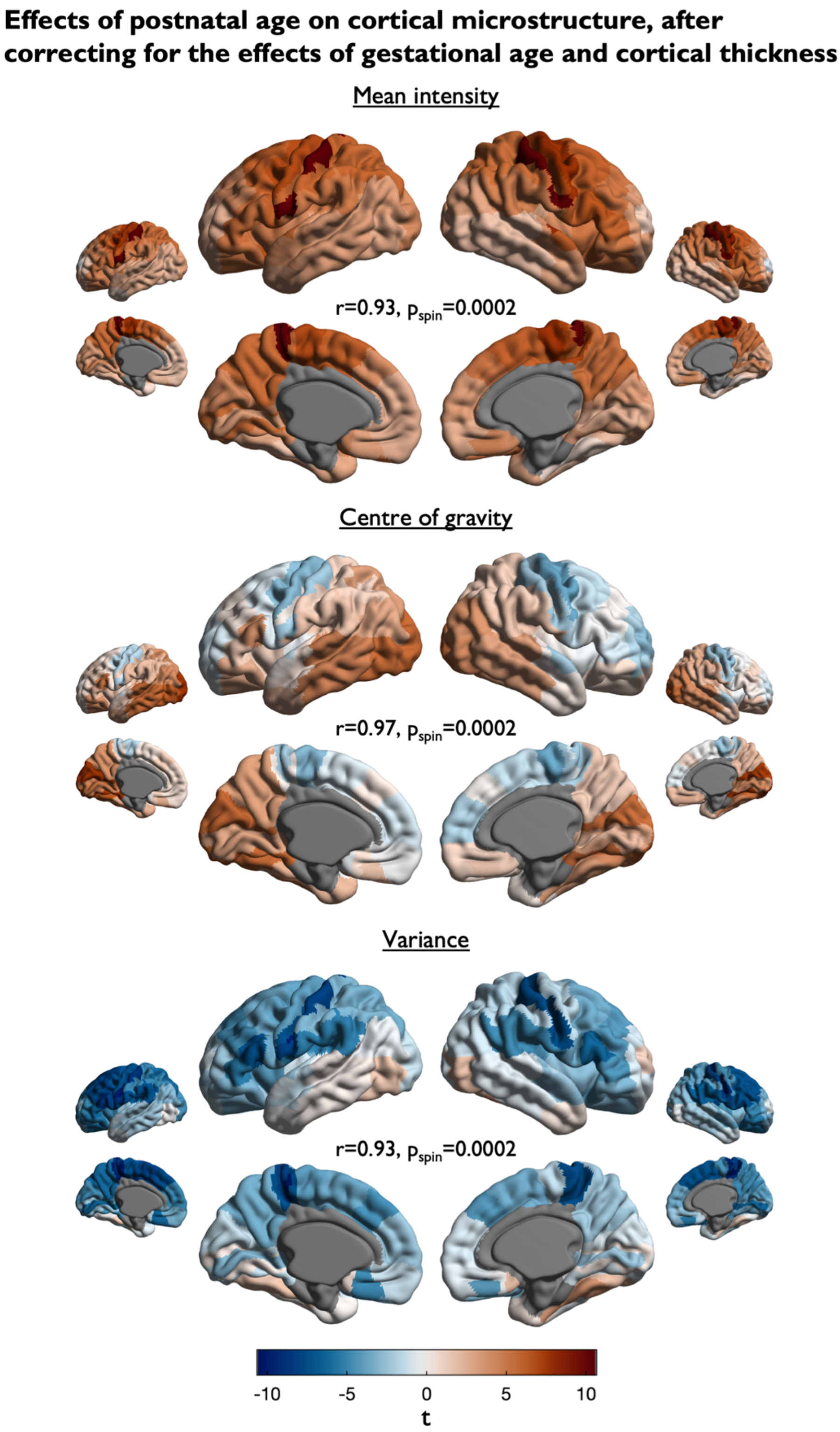


**Supplementary figure 9:** Linear regression models were used to assess the association between postnatal age (PNA) and intracortical profile moments across cortical regions, while controlling for cortical thickness and sex. The original results of this study, without controlling for cortical thickness are displayed as smaller surface maps. The spatial correlation between the effects with and without controlling for cortical thickness is displayed in the middle of each set, together with the p-value derived from spin-based permutation testing (n = 10000), between the 2.5th to 97.5th percentile of the permuted correlations. Surface maps display t-values for the PNA-estimate, projected onto the cortical surface for mean intensity (top), centre of gravity (middle) and variance (bottom). Excluded parcels are displayed in grey.
